# Trends in plant ecology research in Ethiopia (1969-2019): Systematic analysis

**DOI:** 10.1101/2020.09.28.316463

**Authors:** Kflay Gebrehiwot, Sebsebe Demissew

## Abstract

The objective of this paper was to systematically analyze the trend of plant ecological research in Ethiopia. The inclusion and exclusion of the articles for analysis were performed using Reporting Standards for Systematic Evidence Syntheses (ROSES) flow diagram developed for systematic review/meta-analysis. The number of articles published, authors, and collaboration has increased dramatically since the 1960s. Most of the research largely focused on the Dry evergreen Afromontane Forest and grassland complex (DAF) and Moist evergreen Afromontane Forest (MAF) vegetation types, comprising of about 52.6%. Of the remaining vegetation types, the woodlands (14.3%) i.e. *Acacia-Commiphora* woodland and bushland proper (ACW), and *Combretum-Terminalia* woodland and wooded grassland (CTW), desert and semi-desert scrubland (DSS) (2.3%), and the Afroalpine (AA) and Ericaceous Belt (EB) (1.5%) received little attention. A descriptive study on plant community ecology revolving on floristic composition and community structure is the dominant research theme, which revealed a narrow theme in contrast to the global trend. Other plant ecological studies such as reproductive and dispersal ecology of invasive plant species, and pollination ecology seems largely a neglected topic by the academia. Furthermore, the recommendations forwarded are not result-based. As a future direction, the Ethiopian government should develop a project database both for completed and ongoing projects.

## Introduction

Plant ecology as a standalone discipline of botany has a long history. It is highly linked to the works of Alexander von Humbolt in the early nineteenth century [1]. Subsequently, some branches of plant ecology such as synecology and autecology, which emphasizes on community ecology and individual species respectively, emerged. From the early 19^th^ century onwards, plant ecologists have studied stands of vegetation, which they considered samples of a plant community [2]. Nowadays, however, traditional ecological terms (synecology and autecology) are replaced by specialties such as population ecology, community ecology, ecosystem ecology, ecological modeling, global change biology, and remote sensing [1,3,4].

Even though the debate is continuing in the 21^st^ century, the community ‘discrete’ [5] and community ‘continuum’ [6] were at the center of two different views. Vegetation ecology research which focused on plant communities, the effect of abiotic environment on plant distribution, had been the center of ecological research during the first half of the twentieth century [1]. However, in the late 1960s advances in population genetics and the evolutionary theory shifted the theme of plant ecological research into a broader scope, which includes population ecology that combined mathematical modeling and experimentation to investigate population growth, dispersal, and competition from an explicitly Darwinian perspective [1,3,4,7,8].

Plant ecology research employs a wide array of methods depending on the objectives set. These might vary in terms of spatial and temporal scale, organizational levels such as species, population, community, and ecosystem. Hence, the approaches could be either classical or advanced [9]. Although the use of some classical approaches could be retained, several plant ecology research methods are being evolved. Consequently, various research in plant ecology are becoming more reputable than earlier ones. This has a massive contribution to vegetation management and biodiversity conservation.

According to the information extracted from different sources, plant ecology research in Ethiopia started in the late 1960s. The objective of the, probably, first empirical plant ecology research in Ethiopia was to test the community ecology hypotheses [10]. Since Beals’ publication, significant numbers of plant ecological research have been conducted.

Peters [11] criticized and argued that ecology as science hasn’t grown up rather the science moves forward slowly. On his critical response to Peters’ critics and argument, Grace [3] countered that ecological research is growing both in scope and citation impact as well. However, there is no empirical data on the trend of plant ecology researches in Ethiopia. Furthermore, the progress of plant ecology research in the country is not known to date. Consequently, it is challenging to judge the impact of these researches on policy development and conservation actions. Therefore, a thorough bibliographic analysis of plant ecology research in Ethiopia is inevitable to understand if the discipline is growing up as a science. This could have a crucial role in showing what has been done so far and what should be in this field.

The objective of this paper is thus to systematically analyze the trend of plant ecological research in Ethiopia with the aim to provide responses to the questions: i) what are the most researched vegetation (natural ecosystems such as forests) and land use types (the purpose for which the land is used such as farmland) ii) What are the most researched domains of plant ecology, iii) are plant ecology researches influencing national policy, and iv) are there financial resources to fund ecological research from government and other sources?

## Methodology

### Data sources and key search terms used

The various research components were filtered and executed from different sources between the years 1969-2019. The sources include Scopus, PubMed, African Journal Online (AJOL), Addis Ababa University (AAU) theses and dissertation repository, and google scholar. The search terms used in executing the articles included from the title, abstract and keywords. The terms included in searching were; “Floristic” AND “Ethiopia”; “woody” AND “diversity” OR “structur” AND “Ethiopia”; “vegetation” AND “ecology” AND “Ethiopia”; “plant” AND “communit” AND “Ethiopia”; “ordination” AND “classification” AND “Ethiopia”; “species” AND “distribution” OR “Model” AND “Ethiopia”; “Restoration” AND “Ecologi”; and “Elevation” OR “Altitud” AND “gradient” OR “Environment” AND “Ethiopia”.

609 articles from Scopus, 21 Ph.D. and MSc theses from Addis Ababa University (AAU) Institutional repository/Electronic Theses and Dissertations, and 20 articles from AJOL were filtered. The theses and dissertations from AAU and articles extracted from AJOL were all used for the analysis. About 56 journals were involved in the systematic review (S1 & 2 appendix). However, the articles from Scopus had to pass through a thorough selection procedure. In the first phase, books and conference papers were excluded. In the second round, articles that emphasize on land use/land cover change though their title includes terms like forest/vegetation cover were excluded. The inclusion and exclusion of the articles for analysis are shown by using Reporting Standards for Systematic Evidence Syntheses (ROSES) flow diagram developed for systematic review/meta-analysis [12] (Fig 1).

**Fig 1.**
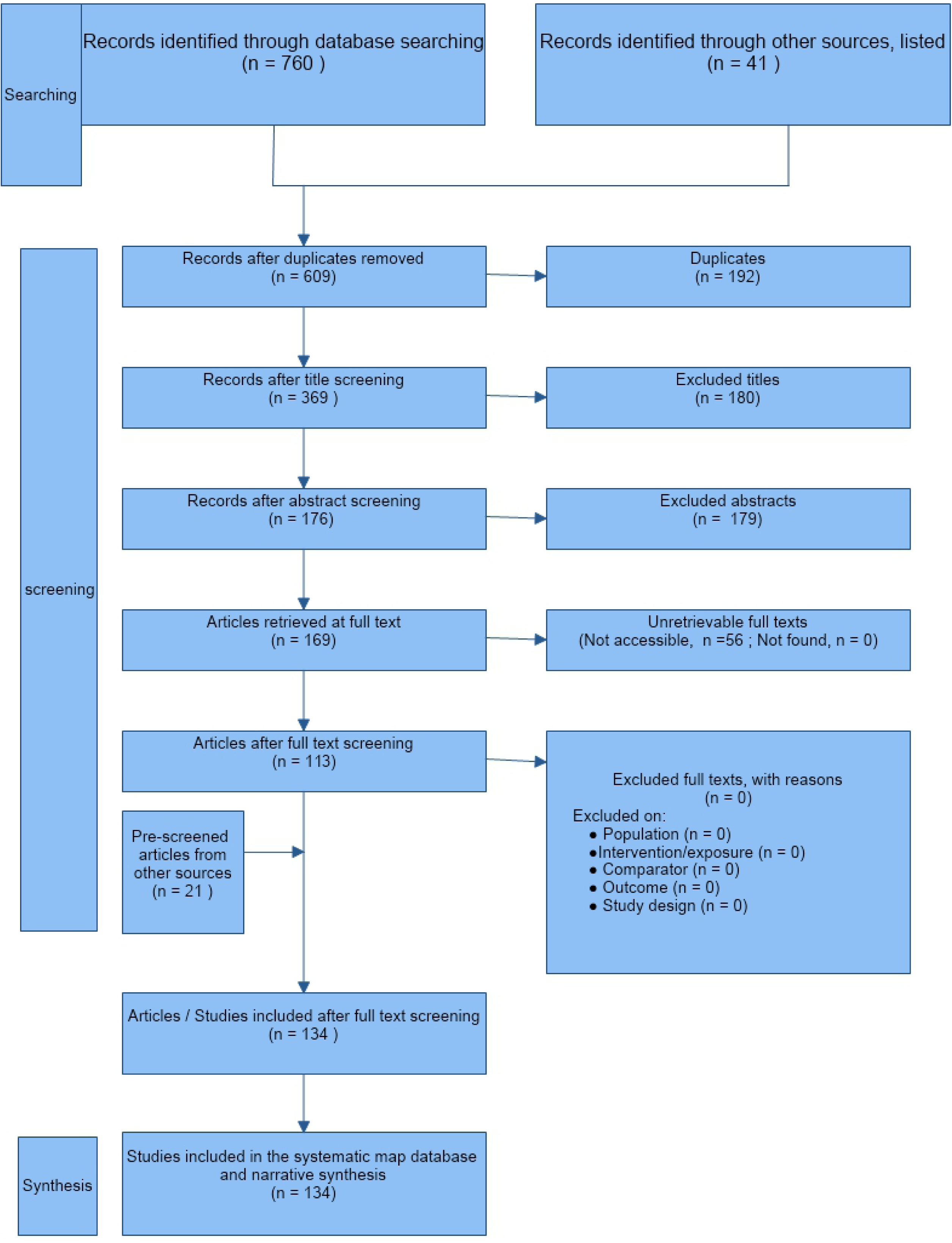
ROSES flow diagram for inclusion and inclusion of research articles from several databases. Modified from Haddaway et al. [12]

**Fig 1.**
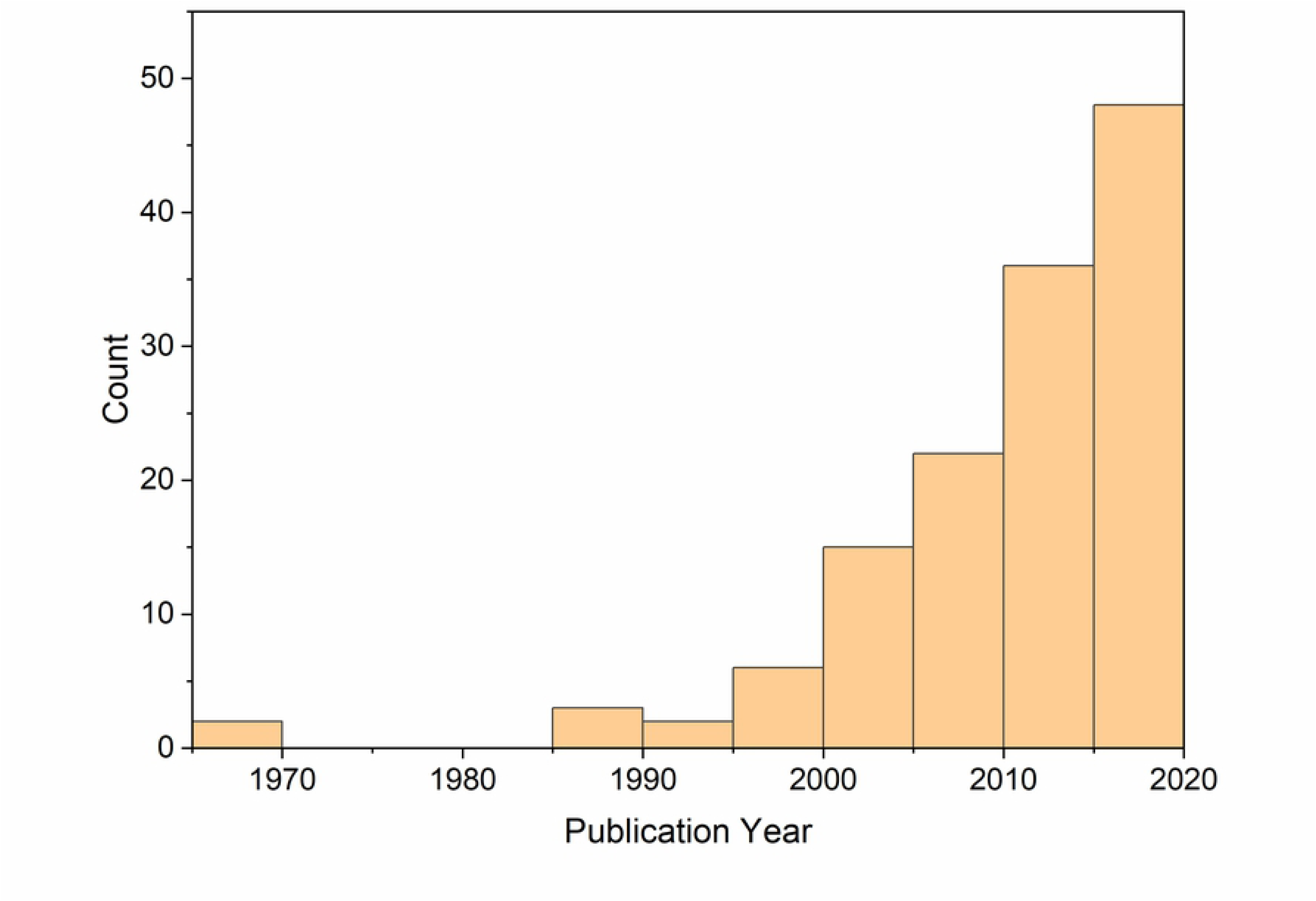
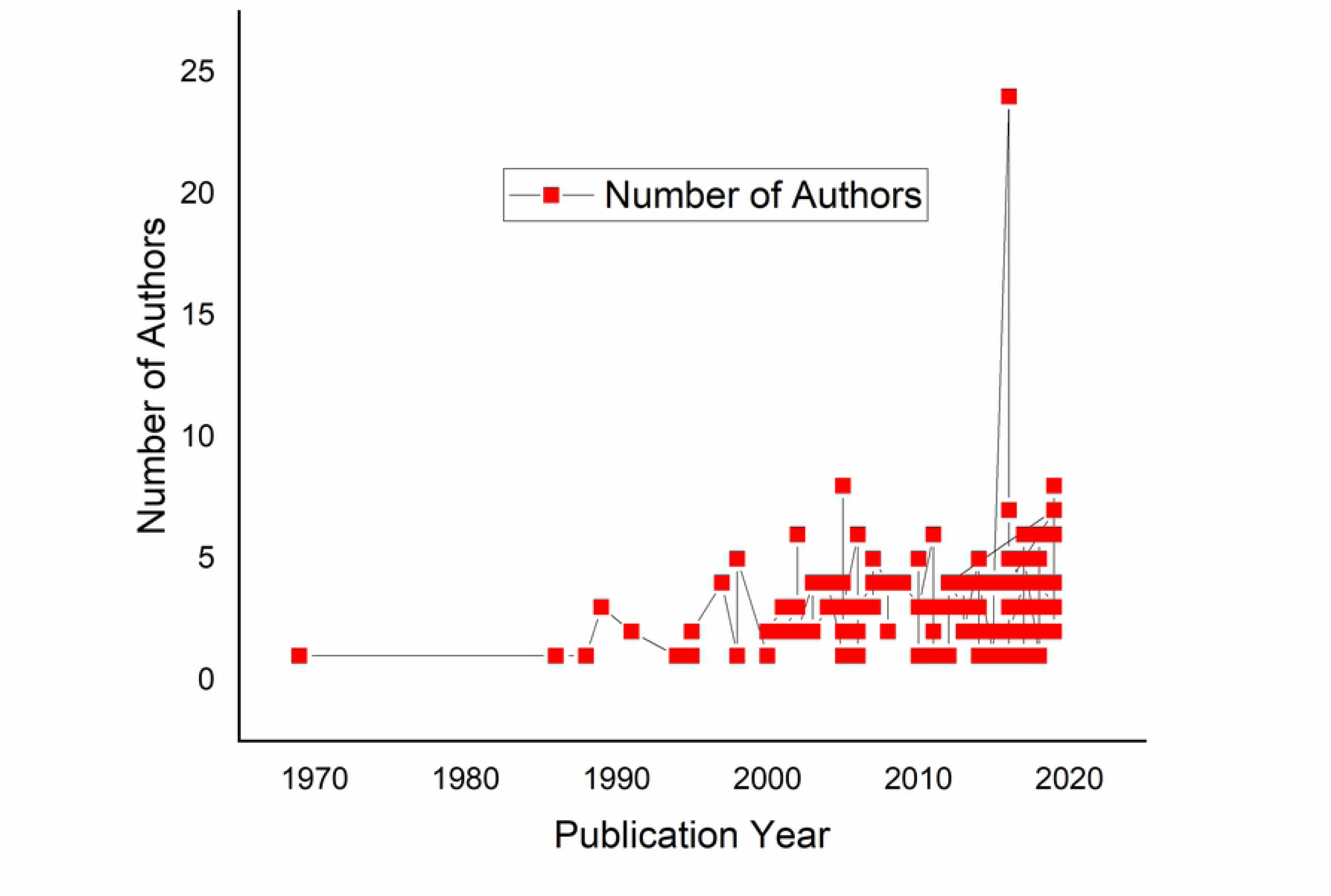
Trend of plant ecology research in Ethiopia. (a) Number of published articles and (b) trend of plant ecological research collaboration

### Data analysis

Pre-analysis coding system for the variables was performed (Table 1). The Authorship and collaboration, plant ecological research components of research, descriptive/experimental, vegetation types, community types, methods employed (sampling and analysis), recommendations, and funding were coded. Descriptive statistics were employed for the analysis.

**Table 1.**
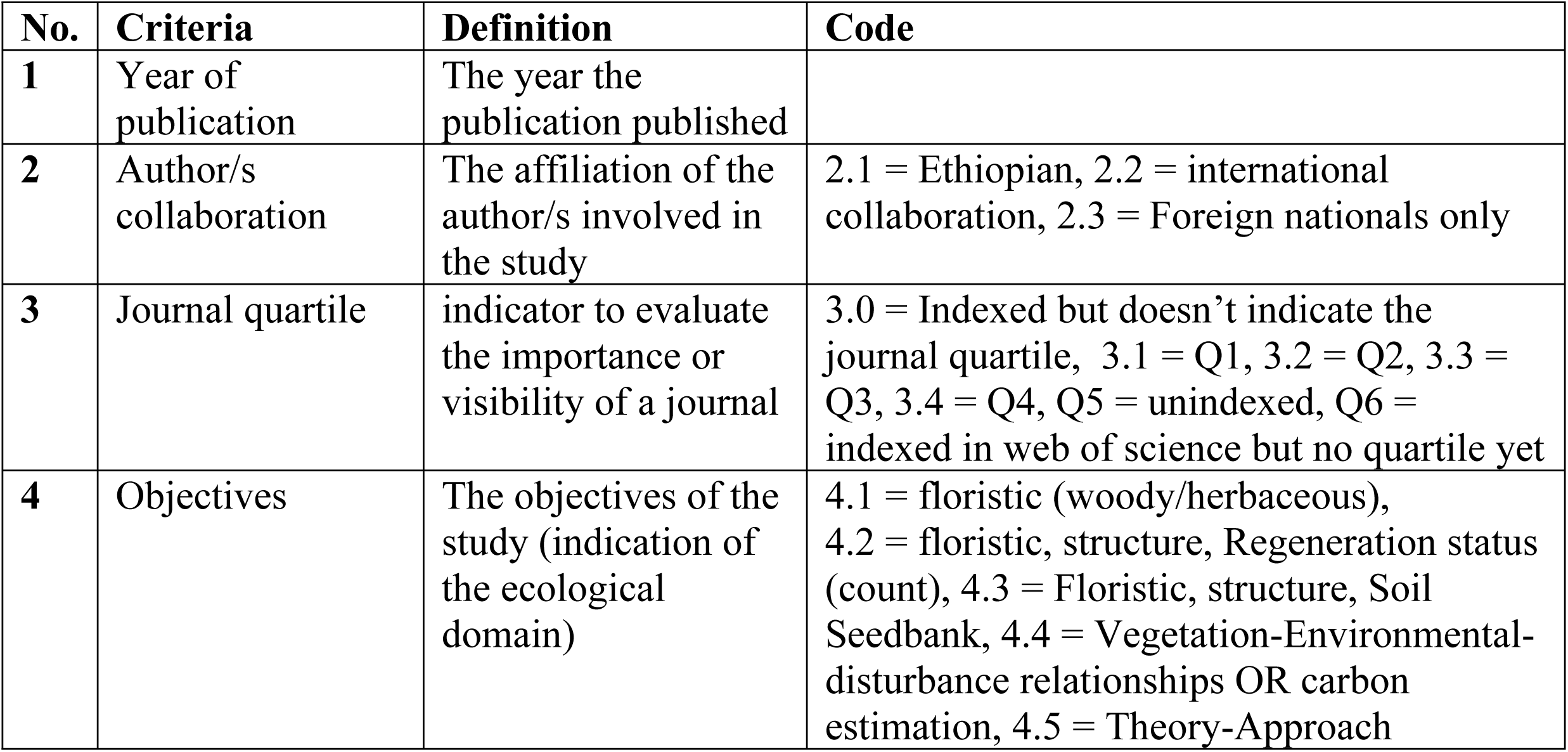
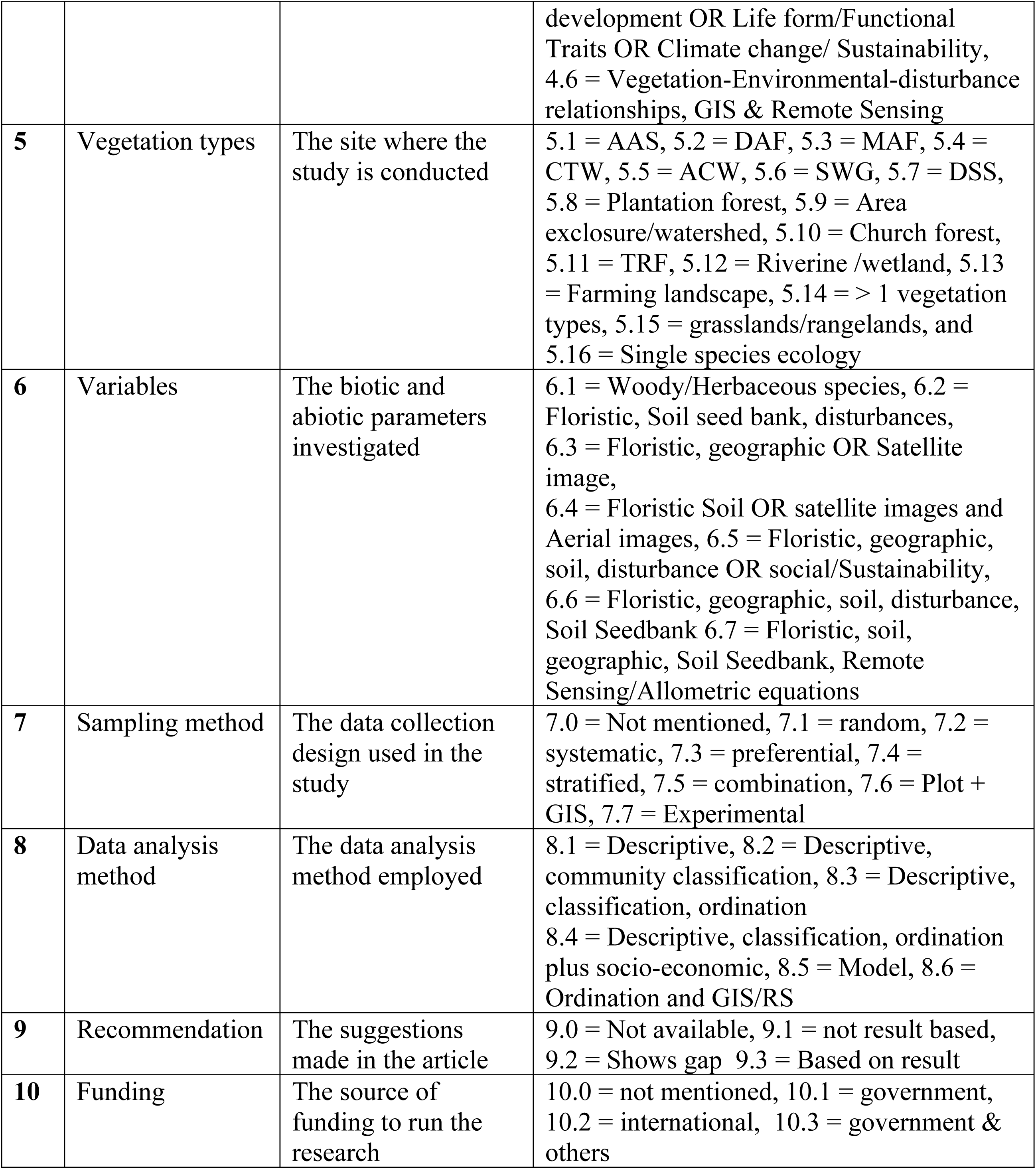
Pre-analysis coding system for all category

## Results

### Authorship and collaboration

Vegetation ecology research in Ethiopia started in the late 1960s (Fig 2a). However, there have been interruptions beween 1960s and 1990s. There has been a dramatic increase after the 1990s. The minimum and maximum articles published in 1969 and 2018 were 2 and 27 respectively. Similarly, the number of authors also showed an increasing trend (Fig 2b). The highest number of authors, 24, was recorded in 2018.

**Fig 2.**
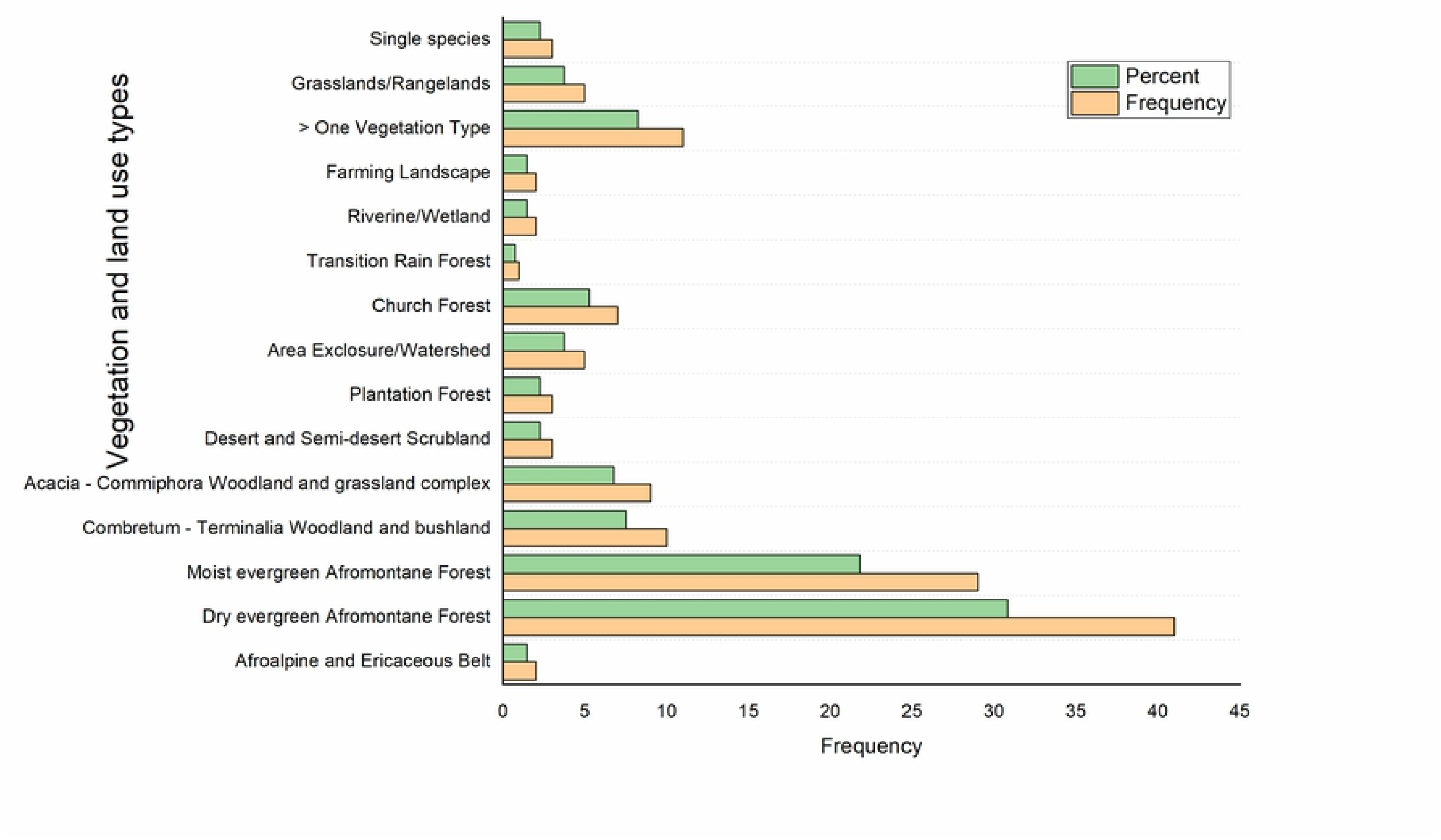
Distribution of the studies in vegetation types and land uses

The author/s affiliation/collaboration involved showed local authors are dominant, 83 (62%). On the other hand, international collaborators and foreign authors only comprised 48 (35.8%) and 3 (2.2%) respectively. International collaborations had started in the late 1990s and grew steadily. On the contrary, the research conducted by foreign nationals only is negligible. The only articles published solely by the foreign nationals were in 1969 and 2017.

### Plant ecological research components

Results revealed that most of the articles’ objectives were descriptive (Table 2). A few articles dealt with some advanced objectives. About 42 (31%) of the articles’ objectives were on the floristic survey, community structure analysis, and assessing the regeneration status of a forest based on a seedling, sapling, and mature tree count. Whereas, the Theory-Approach development OR Life form/Functional Traits OR Climate change/ Sustainability theme covered only 5 (3.7%) of the articles. Even though few articles are available about invasive plant species distribution, and economic impact, studies on reproductive and dispersal ecology of invasive species is almost non-existant. Furthermore, pollination ecology seems the neglected topic in the Ethiopian academia. Exclusive to the very few articles on crop pollination such as Coffee (*Coffea arabica*) by honeybee (*Apis mellifera*), pollination ecology on indigenous and wild plants is badly missing.

**Table 2.** The thematic research topics in the articles

| Objectives | Frequency | Percent |
| --- | --- | --- |
| Floristic (woody/herbaceous) | 16 | 11.9 |
| Floristic, structure, Regeneration (count) | 42 | 31.3 |
| Floristic, structure, SSB | 28 | 20.9 |
| Vegetation-Environmental-disturbance relationships OR carbon estimation | 35 | 26.1 |
| Theory-Approach development OR Life form/Functional Traits OR Climate change/ Sustainability | 5 | 3.7 |
| Vegetation-Environmental-disturbance relationships, GIS & RS | 8 | 6.0 |
| Total | 134 | 100.0 |

### Descriptive/Experimental

Out of the 134 articles reviewed, only two articles were experimental while the remaining are descriptive. One of these articles investigated the tree regeneration potential of only four species namely *Juniperus procera, Ekebergia capensis, Prunus africana* and *Olea europaea* subsp. *cuspidata* under three conditions i.e., along the interior and edge forest gradients, canopy cover, and grazing intensity [13]. The other article determined the floristic composition and soil seed bank richness using manure and livestock grazing as treatments [14]. The objective of these two articles is in a similar domain.

### Plant ecological research on vegetation and land use types

Most of the the plant ecological research articles in Ethiopia focused on the DAF and MAF vegetation types (Figs 3 & 4). Research on these vegetation types comprised about 52.6%. However, the woodlands (14.3%) i.e. *Acacia-Commiphora* woodland and bushland proper (ACB), and *Combretum-Terminalia* woodland and wooded grassland (CTW), desert and semi-desert scrubland (DSS) (2.3%), and the threatened Afroalpine (AA) and Ericaceous Belt (EB)(1.5%) received little attention. The Transitional Rain Forest (TRF) vegetation type was represented by only one article. Furthermore, nearly 8.3% of the studies covered more than one vegetation types. The church forests, grasslands/rangelands, area exclosures comprised 5.3%, 3.8% and 3.8% respectively. Apart from the natural vegetation types, other land uses have also been an area plant ecological research. Plant ecology research on farmland landscape and plantation forests comprised 1.5 and 2.3% respectively. About 2.3% of the studies were focused on single species. Nevertheless, the absence of appropriate geographical coordinates make tracing some of the study sites extremely challenging.

**Fig 3.**
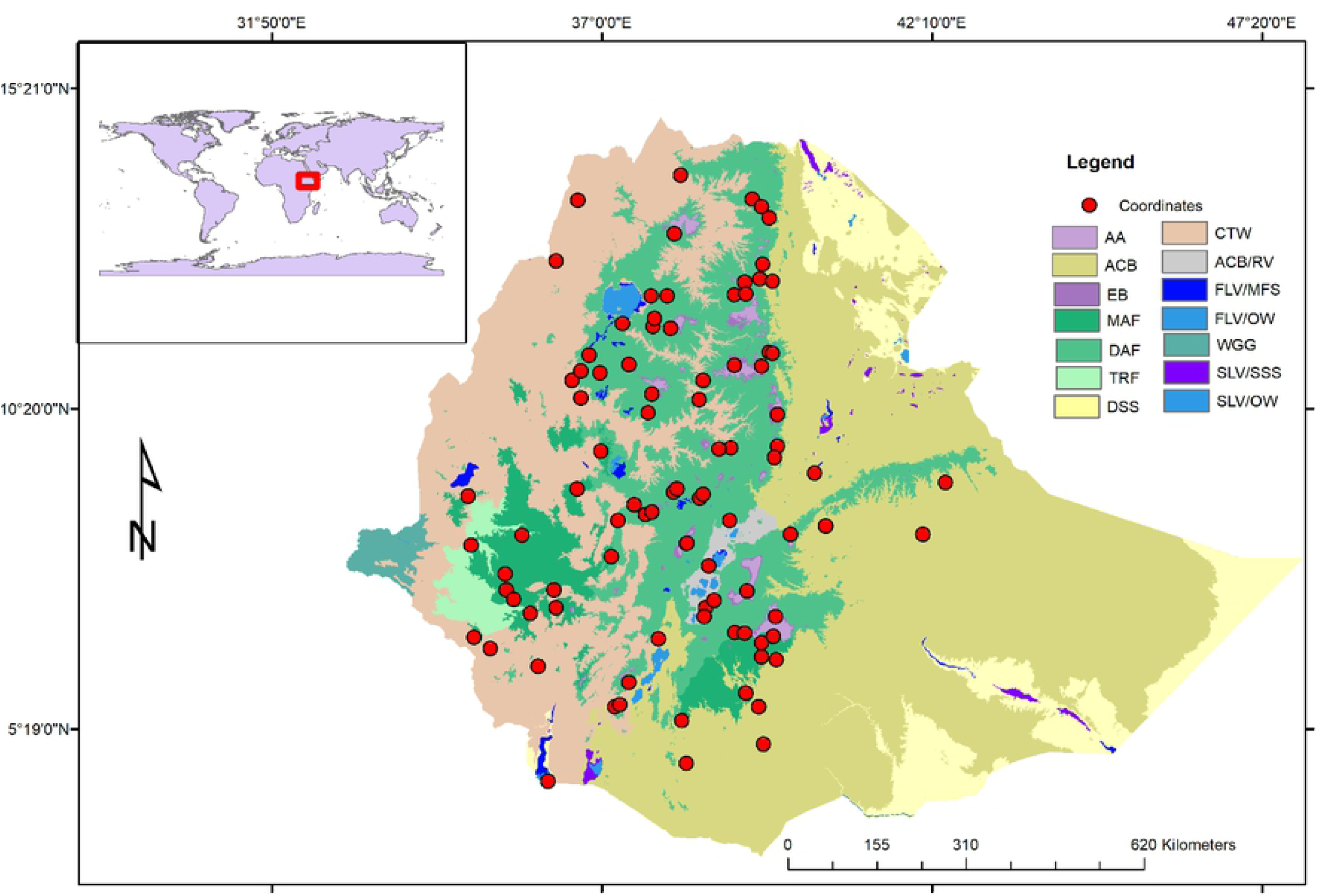
Map of the studies with formal coordinates (red dots) and the corresponding vegetation types (AA = Afroalpine Belt, ACB = *Acacia-Commiphora* woodland and bushland proper, EB = Ericaceous Belt, MAF = Moist evergreen Afromontane Forest, DAF = Dry evergreen Afromontane Forest, TRF = Transitional Rain Forest, DSS = Desert and semi-desert scrubland, CTW = *Combretum-Terminalia* woodland and wooded grassland, ACB/RV = *Acacia* wooded grassland of the Rift Valley, WGG = Wooded grassland of the Western Gambela region, FLV/MFS = Freshwater marshes and swamps, floodplains and lake shore vegetation, FLV/OW = Freshwater lakes - open water vegetation, SLV/SSS = Salt pans, saline/brackish and intermittent wetlands and salt-lake shore vegetation, and SLV/OW = Salt lakes - open water vegetation) [15,16].

**Fig 4.**
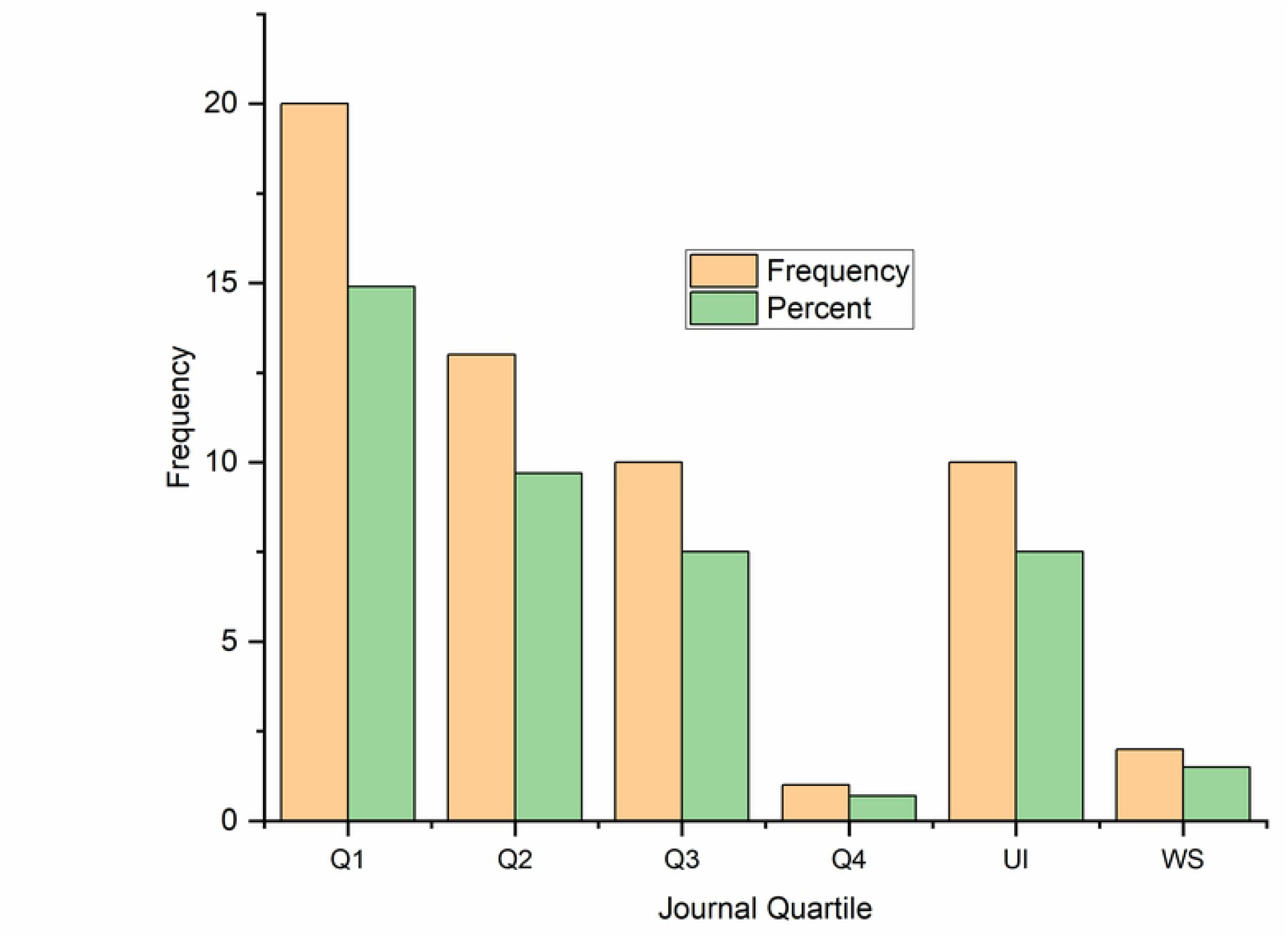
Journal quartile of the articles reviewed (Q1 = quartile 1, Q2 = quartile 2, Q3 = quartile 3, Q4 = quartile 4, UI = Unindexed, WS = indexed in web of Science but no quartile value yet). The Journal quartile value is obtained from Scimago (https://www.scimagojr.com/)

### Plant Community types

About 43% of the studies reported plant community types analysis. The number of communities varied from 2 to 9 with a mean of 5 communities. However, some articles didn’t follow the standard community (*Juniperus – Olea* community) naming procedure while others showed some deviation in the characteristic species of vegetation types. For example, *Olinia rochetiana* (Oliniaceae) is described as a characteristic species of the DAF [15]. However, this species was reported as a characteristic species of MAF. Furthermore, although *Erica arborea* is a characteristic species of Ericaceous Belt, about four articles reported it as a community of the DAF. Furthermore, a shrub/tree and herb (for example, *Albizia schimperiana - Hypoestes forskaolii, Hyparrhenia filipendula - Combretum molle*) were frequently used to name a plant community. Others also named plant community after a weed such as *Achyrantes aspera*. Overlap of plant community types between vegetation types were also reported in different articles. For example, *Arundinaria alpina* and *Maesa lanceolata - Brucea antidysenterica* communities were reported both from DAF and MAF.

### Methods employed by the studies

The selection of data collection and analysis methods might be based on the availability of time, fund, expertise, and objectives amongst others. In the present review 86 (64.2%) of the studies employed systematic sampling while a combination of sampling methods and plot-based data collection supported by GIS only accounted 6 (4.4%) studies. The analysis methods also revealed above 50% of the articles were descriptive (Table 3).

**Table 3.**
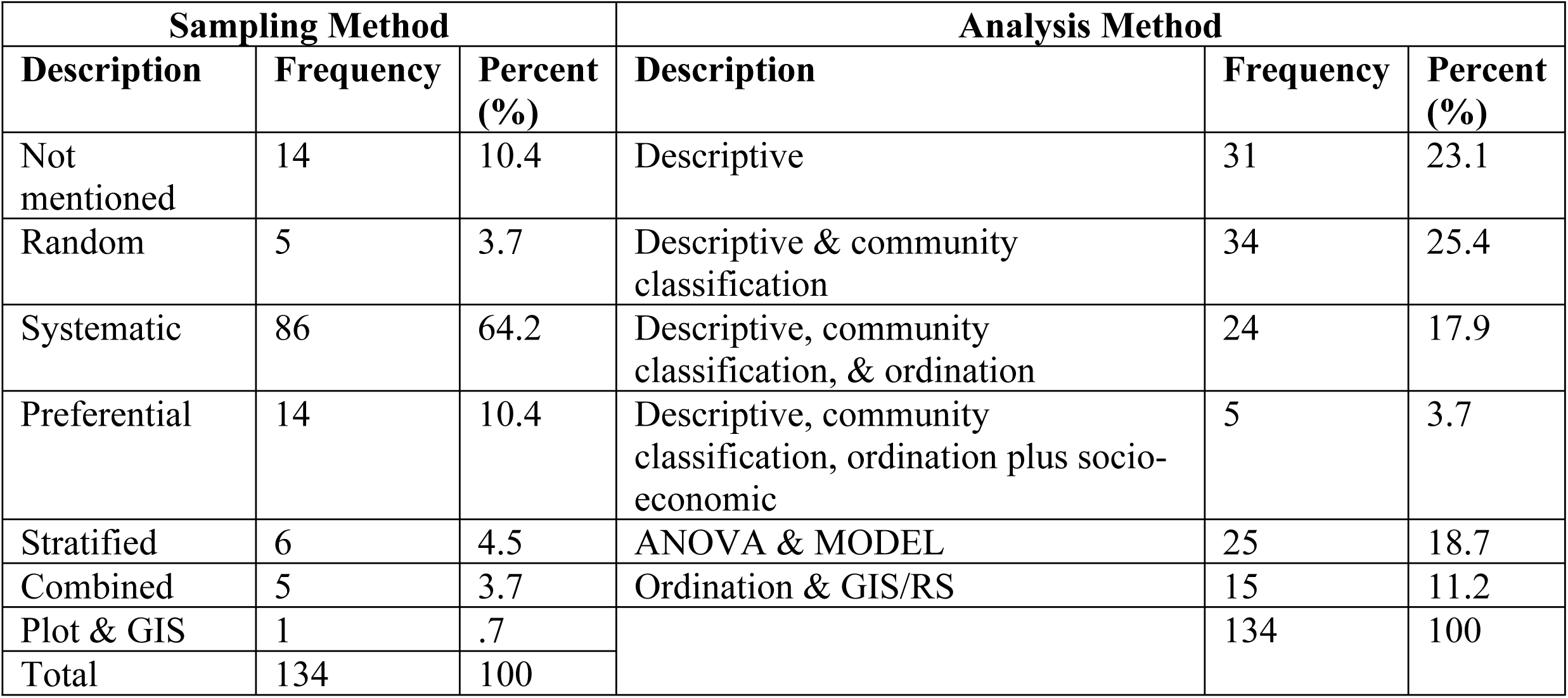
Descriptive statistics of the sampling and analysis method of the studies reviewed

| Sampling Method |  |  | Analysis Method |  |  |
| --- | --- | --- | --- | --- | --- |
| Description | Frequency | Percent (%) | Description | Frequency | Percent (%) |
| Not mentioned | 14 | 10.4 | Descriptive | 31 | 23.1 |
| Random | 5 | 3.7 | Descriptive & community classification | 34 | 25.4 |
| Systematic | 86 | 64.2 | Descriptive, community classification, & ordination | 24 | 17.9 |
| Preferential | 14 | 10.4 | Descriptive, community classification, ordination plus socio-economic | 5 | 3.7 |
| Stratified | 6 | 4.5 | ANOVA & MODEL | 25 | 18.7 |
| Combined | 5 | 3.7 | Ordination & GIS/RS | 15 | 11.2 |
| Plot & GIS | 1 | .7 |  | 134 | 100 |
| Total | 134 | 100 |  |  |  |

### Journal quartile of the articles published

In terms of journal quartile, most of the articles (24.6%) were published in the first quartile and second quartile (Fig 5). The articles that were retrieved from African Journals Online (AJOL) and Addis Ababa Dissertation/Theses repository were not indexed though Momona Ethiopian Journal of Science (MEJS) is indexed in Web of Science and tracked for impact.

### Recommendations forwarded by the studies

Most of the studies’ recommendations are not based on the results of the research (38.1%, n = 51). For example, some floristic composition studies recommended establishment of ‘Natural reserve’ or ‘Biosphere Reserve’. Although floristic study is part of biosphere reserve, proposing ‘Natural reserve’ or ‘Biosphere Reserve’ based on solely floristic list is far beyond the minimum requirement.Only 21.6% (n = 29) of the results revealed that the recommendations are based on the results. On the other hand, 16.4% (n = 22) of the recommendations showed the gaps that were not covered in their studies. On the other hand, several studies (23.9%, n = 32) did not imply any recommendation.

### Funding source reported

The funding source for the articles reviewed was dominantly from international funders 43.3% (n = 58) and government 35.1% (n= 47). The remaining articles’ funding sources were either collaborative (8.2%, n = 11) or did not mention (13.4%, n = 18) their funder. Although the international funding sources have made a significant contribution, the thematic of most of the articles was descriptive as shown in the previous sections.

## Discussion

### Authorship and collaboration

The very few plant ecological studies in the late 1960s are associated to the lack of trained manpower and political instabilities in the country. Exclusive to Haramaya and Gondar, which were colleges of agriculture and health science respectively, there has been only one university with natural science programme, the so-called Haile Selassie I University, current Addis Ababa University since the early 1970s. Furthermore, most of the experts were foreigners with limited knowledge about the vegetation of Ethiopia.

However, from the early 1980s, thanks to the flora project funded by Swedish International Development Cooperation Agency (SIDA), a Swedish organization, experts were trained on several disciplines such as plant ecology, plant systematics, ethnobotany, etc. Few of them are still staff members of Addis Ababa University. Furthermore, with the number of Universities in the country increasing from one in the 1960s to nearly 50 after fifty years. Concurrently, the number of publications rise steeply in the last five decades.

Globally, the extinction of single-authored articles has been confirmed in various disciplines [17]. Similarly, in the present study, single-authored articles showed a declining trend. This is mainly due to the fact that the research projects are either multidisciplinary or students and staff resulting with multiple authorship articles produce better quality research. Furthermore, most of the research are theses/dissertation outputs which include theses/dissertation supervisors and students contribuiting to multi-authored articles.

### Themes of the research topics

In the 21^st^ century, researches are supported by many technology products such as equipment for experiments, software programs for statistical analysis, and other applications such as GIS and remote sensing which support field works. Consequently, the number of plant ecology articles published and the research theme is dramatically increasing globally [18]. In the present systematic review, however, the research theme is very narrow in contrast to the global trend. McCallen et al. [18] identified nearly 50 thematic areas in ecological research. However, it is less than five thematic areas, which were covered in the reviewed articles. A descriptive study on plant community ecology revolving on floristic composition and community structure dominated the studies. The dominance of community ecology research was reported by Réale [19] in their review based on co-citation analysis although it was in the 1970s. This is, however, in contrast to a report by Carmel et al. [20] in which single species research found to be dominant.

Ethiopia’s natural environment is extremely degraded. As a result, most of the environment in the country is a fertile ground to invasive alien, particularly, plant species. Numerous plant species have been recognized as invasive in Ethiopia. Although most of them are exotic, there are few native plants which turned into invasive. I have observed some of the species in the desert and semi-desert scrubland at elevation lower than 1000 m a.s.l. (e.g. *Parthenium hysterophorus* and *Prosopis juliflora*) and Afromontane forests at elevation above 2500 m a.s.l. These means the protected areas in the lowlands and highlands are prone to invasive species. However, a few studies investigated the distribution and impact of invasive species.

Their impact is profound in the rift valley. However, studies on this topic as well as this ecosystem are extermly limited. The persistent environmental degradation, increasing trend of invasive species, and climate change, is aggravating the impact of invasive species. Hence, research on this field is critically in need. Even though the study on invasive species distribution and cover is extremely important for its management, understanding the reproductive and dispersal ecology of the species also play a cruciale role either for management or eradication of the undesirable invasive species.

Pollination is the core of ecological networks. Pollination, the often mutualistic interaction, plays a vital role in maintaining community stability and ecosystem function. Nevertheless, these interactions are threatened due to natural and anthropogenic disturbances leading to a global decline of pollinators. Plant-pollinator networks can potentially modify the population dynamics and the occupied range of a plant species [21]. However, the contribution of pollination networks as driver of plant distribution and assemblage of plant communities has received little attention. Thus, there is a strong need to characterize plant–pollinator interactions at large spatial scales and especially with respect to dynamic communities, whose compositions and patterns of relative species abundance vary in time and space [21]. Pollinator diversity and abundance plays a crucial role in shaping plant communities.

However, studies revealed that pollinators’ diversity is declining at an alarming though most of the researches are on crop pollinators. In the anthropocene, the globe is warming. This is causing changes in species fundamental and realized environmental niches. As a result, plants are shifting their ranges and phenology. Consequently, this cause a plant-pollinator mismatches which leads to plant and pollinators diversity decline. The most species at risk could be plants that required specialized pollinators while the generalized ones are least affected. This is due to specialized plants will be more prone to pollen limitation because they are less likely to interact with any available pollinator [22]. The impact of low abundance and diversity of pollinators do not only influence individual plant species but do extend to plant communities. Hence, future plant ecology research in Ethiopia should consider pollination ecology.

The limited number of articles from experimental studies and advanced themes could be due to the lack of resources for experimental activities and limited expertise in these fields. Unlike the present study, Carmel et al. [20] and Asselin and Gagnon [4] revealed both observation and experimental studies shared almost equal contributions in their systematic review.

Although the Swedish organization (SIDA) made a contribution to training experts from different fields, the theme were bound to basic sciences. Hence, these experts have worked on descriptive research, and the encourage the students they supervised them to work their theses/dissertation on similar topics. Once, the graduates are distributed to different universities, they follow the footsteps of their supervisors. As a result, redundancy of research thematic occurs throughout the country.

### Vegetation and plant community types

Performing ecological research is not an easy task. Taking ecological data from the field requires physical fitness, withstanding harsh conditions, and other field phenomena. Most of the plant ecological research was in either the DAF or MAF vegetation types. This could be due to two main reasons. First, the highest priorities were given to forests and most of the forest cover in Ethiopia is found in these vegetation types. Second, the DAF and MAF are found in a relatively suitable climate where most of the population is currently residing. Furthermore, they are easily accessible. As a result, these are the most threatened vegetation types [23] which needs empirical research.

However, the other important vegetation types attracted little attention. Particularly, the Afroalpine and Ericaceous belt, which are the main sources of freshwater for the downstream population unexpectedly, received little attention. These could be due to either the unwelcoming environmental conditions such as extreme cold and inaccessibility or the little priority given to these vegetation types. Similarly, articles from the vegetation types from drylands/lowlands contributed little.

### Journal quartile of the articles published

Journal impact factor (JIF) quartile is among the widely used indicator to evaluate the importance or visibility of a journal in its field [24]. JIF Q1 means a journal’s impact factor is within the top 25% of the JIF distribution of a particular category showing high impact. On the other hand, Q4 means it is within the lowest 25% of the JIF distribution means lower impact. Hence, the reviewed studies dominantly published their article in relatively high impact journals. Nevertheless, the significant number of studies published in unindexed journals is bleak to trace their impact.

### Recommendation and source of funding

Research recommendations are always for stakeholders who are usually natural resource managers, decision, and policymakers. Thus, it should be based on the empirical data and conclusions made. This helps the stakeholders to decide based on the concrete data and recommendations. Furthermore, showing the thematic research gap would also have profound benefits. However, in the present study, most of the recommendations were not made based on the result. This in some way, shows either missed opportunity to help the stakeholders or obscure in the research objective. However, it doesn’t mean that a study should always wind up by recommendation.

Whether the funding source is government or international, the funders might have their own interest. Oftentimes, funders are interested in making conservation decisions based on scientific evidence [25], to maximize the beneficial outcomes of conservation, given the limited resources available [26]. It is interesting that the international funding dominated in the studies reviewed. However, the research components investigated by using international funding are more or less similar to the investigations conducted by government funding.

Furthermore, monitoring of the completed projects and incorporating the research recommendations into policy and decision-making is poorly noticed. For example, Teketay & Bekele [27], Hundera & Deboch [28] and Hundera et al. [29] recommended Wof-Washa, Gurra Farda, and Dodola forest as a nature reserve. However, they are not designated as a protected area yet [30]. Furthermore, forest and woodland cover loss are reported even in protected areas [31,32], revealing loose policy and decision making. These could be due to three reasons. These are i) the government is responsible for integrating the research outputs into policies, ii) complexity of socio-ecological systems [33], lack of funding for conservation operations [34], and iii) lack of trustworthy scientific studies.

### Is plant ecology research growing up in Ethiopia?

Debates were raised about the terminologies used to describe the development of ecology as a science [3,11]. Grace [3] suggested the question should be ‘is ecology growing up?’, not ‘Has ecology grown-up?’ like Peters [11] asked. Bringing Grace’s question to the Ethiopian perspective, If the frame is the number of published studies, plant ecology research is tremendously growing up. Even though evaluating the quality of research needs a different pathway, the quality of plant ecological in Ethiopia is not growing up. The criteria here is the research theme, methods apllied, and depth of the study. This is excluding the land use/land cover change domain. Plant community, more or less, descriptive studies, dominated the plant ecology research thematic. This topic has been the center of plant ecological research in the 1960s to 1970s in Europe and Northern America. Hence, although it seems lest to bring this thought to the academia, plant ecology in Ethiopia is still in the 1970s.

## Conclusions and future directions

This study systematically synthesized trends of plant ecology research in Ethiopia over the last 50 years. Plant community more or less, descriptive studies, dominated the plant ecology research thematic most of them distributed in the DAF and MAF while the Afroalpine and Ericaceous Belt, woodland (ACB & CTW), and desert and semi-desert scrubland (DSS) vegetation types got little attention. Hence, the following future directions are suggested to improve the forthcoming plant ecology research in Ethiopia.

1. Future plant ecology research should gear to contemporary ecological researches. These include the application of remote sensing in vegetation ecology, climate change and vegetation ecology, plant functional ecology, vegetation temporal dynamics, and experimental approaches.
2. Experimental plant ecological studies are almost non-existant in the documents reviewed. Hence, future research should look into this perspective.
3. Establishing a committee that comprised plant taxonomists, plant ecologists, geologists, geographers, and GIS experts is recommended to investigate and map the plant community types at national level. This is crucial to allocate conservation resources objectively. Otherwise, fragmented studies of plant community types of a particular sites made conservation efforts challenging.
4. Recommendations from any research study, if available, should be based on the empirical information inorder to make policy and decision justifiable.
5. Funders such as government agencies, NGOs, others including international ones are advised to provide resources to cover important topics such as invasive species ecology, application of GIS and remote sensing in vegetation ecology, and community interactions. Supervisors should also take the lion’s share in helping graduate students identify contemporary ecological research areas such as pollination ecology.
6. The Ethiopian Ministry of Science and Higher Education (MoSHE) should establish a database both for completed and ongoing research projects i.e., project registration in addition to what exists at various universities. This would help to avoid project redundancy.

## Supporting information

**S1 Appendix**. Journals included in the review.

**S2 Appendix**. Studies included in the systematic review.

## Reference

1. Hagen JB. History of Plant Ecology. Encyclopedia of Life Sciences (ELS). John Wiley & Sons, Ltd; 2010. pp. 1–5. doi: 10.1002/9780470015902.a0003288.pub2

2. Mueller-Dombois D, Ellenberg H. Aims and methods of vegetation ecology. Toronto, Canada: John Wiley & Sons; 1974.

3. Grace J. Has ecology grown up? Plant Ecol Divers. Taylor & Francis; 2019;12: 387–405. doi: 10.1080/17550874.2019.1638464

4. Asselin H, Gagnon D. Trends in ecological research : reflecting on 21 years of Écoscience. Écoscience. Taylor & Francis; 2015;22: 1–5. doi: 10.1080/11956860.2015.1120511

5. Clements F. Plant Succession: An Analysis of the Development of Vegetation. Carnegie Institution of Washington; 1916.

6. Gleason H. The Individualistic Concept of the Plant Association. Society. 1926;53: 7–26. doi: 10.1007/s

7. Harper JL. A Darwinian approach to plant ecology. J Ecol. 1967;55: 247–270.

8. Harper JL. Population Biology of Plants. London: Academic Press; 1977.

9. Henderson PA. Ecological Methods. Encyclopedia of Life Sciences (ELS). John Wiley & Sons, Ltd: Chichester; 2012. pp. 1–6. doi: 10.1002/9780470015902.a0003271.pub2

10. Beals EW. Vegetational Change Along Altitudinal Gradients: Studies in Ethiopia show that discreteness of zonation varies with steepness of slope. Sci New Ser. 1969;165: 981–985.

11. Peters RH. A Critique for Ecology. Cambridge, UK: Cambridge University Press; 1991.

12. Haddaway NR, Macura B, Whaley P, Pullin AS. ROSES RepOrting standards for Systematic Evidence Syntheses : pro forma, flow - diagram and descriptive summary of the plan and conduct of environmental systematic reviews and systematic maps. Environ Evid. BioMed Central; 2018;7: 7. doi: 10.1186/s13750-018-0121-7

13. Wassie A, Sterck FJ, Teketay D, Bongers F. Tree regeneration in Church forests of Ethiopia: Effects of microsites and management. Biotropica. 2009;41: 110–119. doi: 10.1111/j.1744-7429.2008.00449.x

14. Woldu Z, Saleem MAM. Grazing induced biodiversity in the highland ecozone of East Africa. Agric Ecosyst Environ. 2000;79: 43–52.

15. Friis I, Sebsebe Demissew, Breugel P van. Atlas of the Potential Vegetation of Ethiopia. Denmark: The Royal Danish Academy of Sciences and Letters; 2010.

16. van Breugel P, Kindt R, Lillesø JP., Bingham M, Demissew S, Dudley C, et al. Potential Natural Vegetation Map of Eastern Africa (Burundi, Ethiopia, Kenya, Malawi, Rwanda, Tanzania, Uganda and Zambia). Version 2.0. In: Forest & Landscape Denmark and World Agroforestry Centre (ICRAF). 2015.

17. Barlow J, Stephens PA, Bode M, Cadotte MW, Lucas K, Newton E, et al. On the extinction of the single - - authored paper : The causes and consequences of increasingly collaborative applied ecological research. J Appl Ecol. 2018;55: 1–4. doi: 10.1111/1365-2664.13040

18. McCallen E, Knott J, Nunez-mir G, Taylor B, Jo I, Fei S. Trends in ecology : shifts in ecological research themes over the past four decades. Front Ecol Environ. 2019; doi: 10.1002/fee.1993

19. Réale D, Khelfaoui M, Olivier P, Yves M. Mapping the dynamics of research networks in ecology and evolution using co-citation analysis (1975 – 2014). Scientometrics. Springer International Publishing; 2020; doi: 10.1007/s11192-019-03340-4

20. Carmel Y, Kent R, Bar-massada A, Blank L, Liberzon J, Nezer O, et al. Trends in Ecological Research during the Last Three Decades – A Systematic Review. PLoS Med. 2013;8: e59813. doi: 10.1371/journal.pone.0059813

21. Pellissier L, Alvarez N, Guisan A. Pollinators as drivers of plant distribution and assemblage into communities. In: Patiny S, editor. Evolution of plant–pollinator relationships. Cambridge, UK.: Cambridge University Press; 2012.

22. Knight TM, Steets JA, Vamosi JC, Mazer S, Burd M, Campbell DR, et al. Pollen limitation of plant reproduction: ecological and evolutionary causes and consequences. Annu Rev Ecol Evol Syst. 2005;36: 467–97.

23. Woldu Z. Forest in the vegetation types of Ethiopia and their status in the geographical context. In: Edwards S, Demissie A, Bekelle Y, Haase G, editors. forest genetic resources-conservation: principles strategies and action. Addis Ababa, Ethiopia; 1999.

24. Liu W, Hu G, Gu M. The probability of publishing in first-quartile journals. Scientometrics. Springer Netherlands; 2015; doi: 10.1007/s11192-015-1821-1

25. Burivalova Z, Miteva D, Salafsky N, Butler RA, Wilcove DS. Evidence Types and Trends in Tropical Forest Conservation Literature. Trends Ecol Evol. Elsevier Ltd; 2019;34: 669–679. doi: 10.1016/j.tree.2019.03.002

26. Game E. Cross-discipline evidence principles for sustainability policy. Nat Sustain. 2018;1: 452–454.

27. Teketay D, Bekele T. Floristic composition of Wof-Washa natural forest, Central Ethiopia : implications for the conservation of biodiversity. Feddes Repert. 1995;106: 127–147.

28. Hundera K, Deboch B. Woody Species Composition and Structure of the Gurra Farda Forest, Snnpr, South Wastern Ethiopia. Ethiop J Educ Sci. 2008;3.

29. Hundera K, Bekele T, Kelbessa E. Floristics and phytogeographic synopsis of a dry afromontane confirous forest in the Bale mountain (Ethiopia): implications to biodiversity conservation. Sinet Ethiop J Sci. 2007;30: 1–12.

30. UNEP-WCMC. World database on protected areas user manual 1.5. Technical report. Cambridge, UK; 2017.

31. Nune S, Soromessa T, Teketay D. Land Use and Land Cover Change in the Bale Mountain Eco-Region of Ethiopia during 1985 to 2015. Land. 2016;5: 41. doi: 10.3390/land5040041

32. Arafaine Z, Asefa A. Dynamics of Land Use and Land Cover in the Kafta-Sheraro National Park, NW Ethiopia : Patterns, Causes and Management Implications. Momona Ethiop J Sci. 2019;11: 239–257. doi: http://dx.doi.org/10.4314/mejs.v11i2.5

33. Miteva DA. Evaluation of biodiversity policy instruments: what works and what doesn’t? Oxford Rev Econ Policy. 2012;28: 69–92.

34. Ferraro PJ, Pattanayak S. Money for nothing? A call for empirical evaluation of biodiversity conservation investments. PLoS Biol. 2006;4: 0482–0488.

